# Dynamic strategic social learning in nest-building zebra finches and its generalisability

**DOI:** 10.1101/2025.06.06.658225

**Authors:** Alexis J. Breen, Tristan Eckersley, Andrés Camacho-Alpízar, Connor T. Lambert, Gopika Balasubramanian, Susan D. Healy, Richard McElreath, Lauren M. Guillette

**Affiliations:** Department of Human Behavior, Ecology and Culture, Max Planck Institute for Evolutionary Anthropology; Leipzig 04103, Germany; School of Biology, University of St Andrews; St Andrews, Fife, KY16 9ST, United Kingdom; Department of Psychology, University of Alberta; Edmonton, Alberta, T6G 2R3, Canada

## Abstract

Animals often balance asocial and social information strategically, adjusting when and from whom they copy based on context. Yet the cognition driving this dynamic—and its broader implications—remains poorly understood. We tested whether zebra finches use a *copy-if-dissatisfied* strategy by manipulating the quality of their initial nest-building or reproductive experience, showing them a conspecific nest-builder, and tracking subsequent material choices. Builder-males were more likely to choose the demonstrated ‘social’ material—particularly at first choice—if they had previously used low-quality material. Using cognitive modelling, we estimated how latent learning mechanisms shaped decisions, identifying two asocial and two social parameters. These estimates provide the first formal evidence for the cognitive basis of nest building. Forward simulations informed—but not predetermined—by these parameters approximated observed behaviour, supporting their causal role. We then used these parameters in exploratory simulations to test how choices might shift under novel payoff contexts. We found that payoff structure—not (dis)satisfaction—was the primary driver of social material use, though higher rewards did not proportionally increase copying. These exploratory simulation results offer preliminary insight into mechanisms underlying material-use variation. Our study illustrates how computational modelling can robustly link behaviour to underlying learning mechanisms and probe the generalisability of animal cognition—a rarity in this field.

## 1 Introduction

Strategic social learning refers to a suite of decision rules individuals may use to determine when to rely on socially provided information, such as *copy-if-kin, copy-if-friends*, and *copy-if-dissatisfied* [1]. These strategies require balancing personal experience (asocial information) with public cues (social information), then updating that balance in light of outcomes. Striking this balance is complex: over-reliance on either source could be costly [2]. While behavioural evidence for such strategies is widespread across taxa (recent review: [3]), the cognitive mechanisms that enable this dynamic balancing—especially in non-human animals—remain largely ‘blackboxed’ [4].

By contrast, computational modelling in human studies has made significant progress in ‘unblackboxing’ the dynamic processes underlying strategic social learning [5], from the individual level to the crowd level [6, 7]. This analytical approach typically involves building a statistical model or models to infer cognition from behaviour—quantifying, for instance, the influence of asocial mechanisms like reinforcement learning and social mechanisms like conformity [8]. These mechanistic estimates can then be used in forward simulations, which predict agents’ behaviour based on the inferred cognitive processes [9, 10]. When simulated behaviour aligns with observed data, the model is considered a good approximation of the underlying mechanisms [11, 12, 13]; when it does not, mismatches should prompt transparent reporting of limitations [14]. Such simulations also allow researchers to test how behaviour might change under new contexts, such as altered payoffs [14], to evaluate how well the model generalises beyond the original study [9, 10].

The growing recognition of computational modelling’s explanatory power in non-human animal cognition [15] has likely contributed to its emerging application in studies of strategic social learning in these systems. To date, at least five studies—on wild and temporarily-captive tits [13, 16], wild monkeys [17, 18], and wild chimpanzees [19]—have applied this framework (*sensu* [8]) to infer the roles of asocial and social learning in foraging behaviour. For example, wild tits were found to determine ‘what to eat’ primarily through reward-based individual updating and frequency-based conspecific copying, with these strategies supported by forward simulations [13]. Further such simulations manipulating agents’ copying levels demonstrated that the moderate social conformity observed in wild birds generally maximised foraging returns [13]

Nest building, like foraging, strongly affects birds’ fitness and may favour strategic social learning. Once considered wholly innate, it is now understood to reflect individual, social, and cultural experience [20, 21, 22, 23, 24, 25]. The most comprehensive evidence of experience-dependent changes in avian nest building comes from the Australian zebra finch (*Taeniopygia castanotis*), a colonial species that constructs intricate domed nests. In the wild, zebra finches use stiff grass stems for structure and soft linings such as feathers or wool [26], with densities reaching 76.6 nests per hectare in thickets [27]. After nest predation, they rebuild more than 20 metres away, indicating avoidance learning [28]. They also use social cues: wild individuals tend to nest near conspecifics at similar breeding stages, suggesting synchronisation via prospecting [29, 30]. In laboratory settings, zebra finches build with diverse materials [21], add a dome when material allows, and learn to prefer stiff string that enables more efficient dome construction [31], a behaviour linked to reproductive success [32]. Social learning also guides their material choice: adults prefer the colour of material they observed being handled by adults during their juvenile stage [22], and they abandon their own preferences after seeing an unoccupied nest composed of their non-preferred colour [33]. Critically, zebra finches are selective in the type of social cue they use—copying material choices of familiar, but not unfamiliar, conspecifics [20]. Together, these findings position the zebra finch as a powerful model for investigating strategic asocial and social information use in nest building.

Here, we examined strategic social learning in builder-male zebra finches housed in a laboratory setting (Fig. 1). Specifically, we asked: (i) do males use a strategy supposed to be widespread: *copy-if-dissatisfied* [1]? and (ii) does its use depend on the *type* of dissatisfaction? We induced dissatisfaction in two ways: (1) by having males build with flexible string—a low-quality material that requires greater investment to maximise reproductive success [31, 32]; or (2) by removing clutches during incubation— a manipulation known to influence later material choices [34, 35]. In one group, males built a partial nest using either stiff or flexible string. In another group, all males built with breeder-standard coconut fibre, but only half successfully reproduced due to our experimental clutch removal. We then assessed each male’s initial material preference (orange versus pink string), allowed them to observe a conspecific building with their non-preferred colour, and recorded their choices during a second round of nest building, where both the asocial (preferred) and social (demonstrated) materials were available. If zebra finches use a *copy-if-dissatisfied* strategy, we expected that males given flexible string and/or those whose clutches were removed, would shift their material preference toward the demonstrated social material. We further predicted that dissatisfaction induced by recent reproductive failure would exert a stronger effect than dissatisfaction following building experience, consistent with previous findings [31].

**Figure 1.**
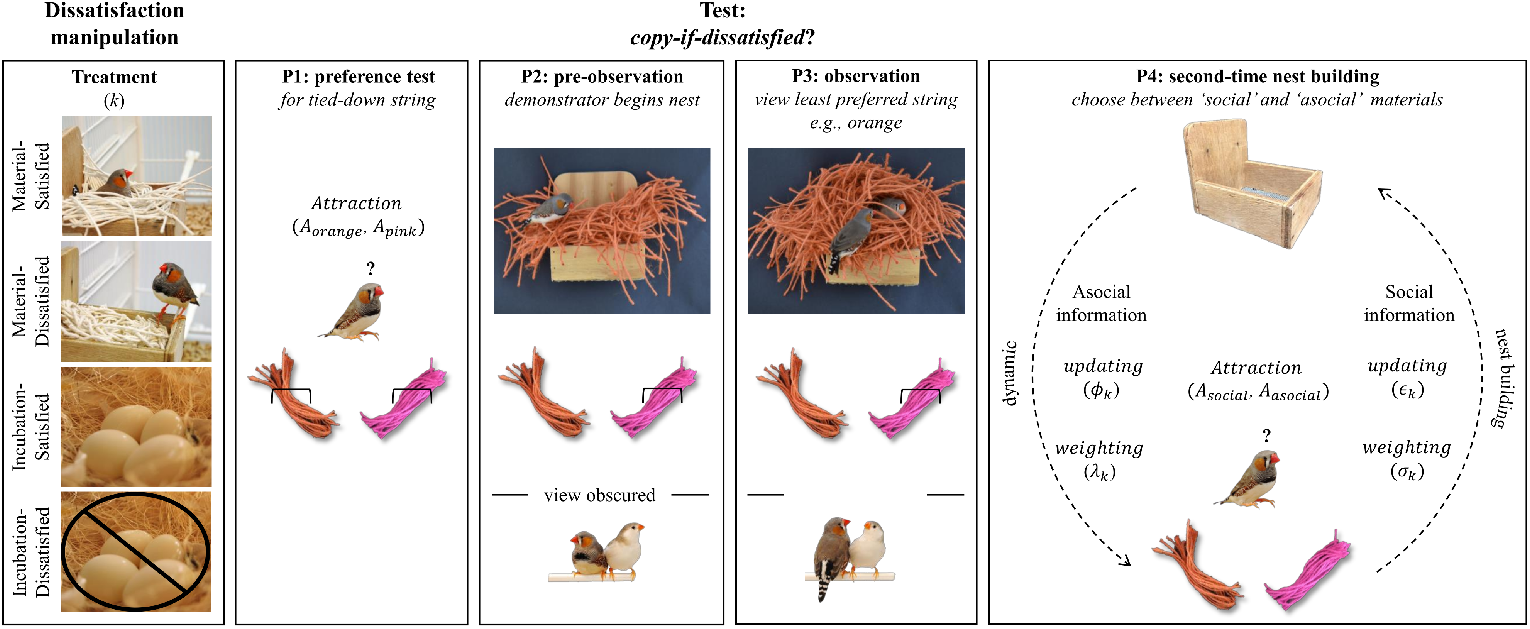
Protocol. The study comprised a Dissatisfaction Manipulation (left column) and a Test with four consecutive phases (P1–P4; columns 2–5) involving subject males and female partners. Dissatisfaction occurred in two contexts: nest building (Material-Satisfied: stiff string; Material-Dissatisfied: flexible string) or incubation (Incubation-Satisfied: eggs hatched; Incubation-Dissatisfied: eggs removed). All males then proceeded to the Test. In P1, each male’s pink versus orange string preference was assessed using tied-down bundles to prevent nest building. In P2, a demonstrator built a nest with 100 pieces of the male’s non-preferred material while visually occluded; the preferred material was present but tied down. In P3, after the demonstrator reached a standardised endpoint, the male observed him continue with the same material. In P4, the male built a second nest, choosing between the now loose demonstrated (social) and non-demonstrated (asocial) materials. The schematic (far right) shows the learning model: attraction values update from asocial (*ϕ, λ*) and social (*ϵ, σ*) information, dynamically influencing nest building. Images: Eira Ihalainen, Patricia Stockebrand, and Animal Cognition Research Group.

These behavioural predictions, however, cannot reveal hidden cognitive mechanisms [10]. To address this, we fit hierarchical Bayesian experience-weighted attraction (EWA) models—originally developed for human strategic behaviour [36]—to infer four latent learning parameters that could jointly underpin material choices:

- *ϕ*, how rapidly males refine material preferences (asocial information updating);
- *λ*, how strongly males exhibit material preferences (asocial information weighting);
- *ϵ*, how elastically males react to deposited social material (social information updating);
- *σ*, how much males consider asocial versus social information (social information weighting).

These estimates allowed us to capture each nest-building male’s decision-making and run forward simulations for two purposes: first, to ensure the model-generated behaviour resembled real data; and second, to test the robustness of our inferred mechanisms in new contexts. In our study, choosing the social material offered no real benefit. But in other contexts, material choice often conveys advantage, such as within-group social signalling (e.g., coloured nest material in birds [24, 37]) or efficiency (e.g., tool use in primates [38, 39, 40]). These choice behaviours may be culturally maintained via social learning [23, 40], and even reflect cumulative culture [39]. To crudely simulate such contexts, we imposed two hypothetical payoff structures: (1) choosing *social pays*, where the social material yielded a payoff and the asocial material none (one versus zero, respectively); and (2) choosing *social pays more*, where the social material yielded a payoff twice that of the asocial material (two versus one, respectively). Thus, we probed how differential (dis)satisfaction influenced material choice under contexts beyond our study where *copy-if-dissatisfied* is advantageous or increasingly so.

## 2 Methods

### 2.1 Subjects

Subjects were 75 male–female adult (*>*90 days) zebra finch pairs. Experiments took place in two laboratories (Experiment 1: Saint Mary’s Animal Unit, University of St Andrews, UK; Experiment 2: Department of Psychology, University of Alberta, Canada). Birds in Experiment 1 were bred in-house; those in Experiment 2 came from a breeder (Eastern Bird Supplies, Quebec, Canada). In both experiments, zebra finches participated as subject pairs (Experiment 1: *n* = 19; Experiment 2: *n* = 34) or demonstrator pairs (Experiment 1: *n* = 16; Experiment 2: *n* = 6). All subject pairs were first-time builders. Further background, housing, and husbandry details are in the *Supplementary material*.

### 2.2 Experimental protocol

In both Experiments 1 and 2 (Fig. 1), we varied the quality of subject males’ first building/incubation experience (the dissatisfaction manipulation). Afterwards, subject pairs were individually transferred, in blocks, into one side of a test cage initially separated from the other side by an opaque, removable barrier. The test comprised four consecutive phases (detailed below).

#### 2.2.1 Dissatisfaction manipulation

In Experiment 1, we manipulated dissatisfaction during nest building by randomly assigning males to build with 100 pieces of either (1) polished ‘stiff’ string or (2) unpolished ‘flexible’ string, forming the *Material-Satisfied* (Mat-Sat) and *Material-Dissatisfied* (Mat-Diss) treatments. All string was off-white, 3 mm in diameter, cut into 15 cm lengths (James Leaver Co., London, United Kingdom; see Figure 1a in [32]) and placed in a single pile at the cage floor centre, aligned with the wall-mounted nestbox. The manipulation ended once males deposited all 100 pieces into the nestbox, after which both nest and nestbox were removed (mean days ± SE to build a partial nest: Mat-Sat 5.00 ± 1.98; Mat-Diss 4.78 ± 1.36). This experience level is sufficient to shape subsequent material preferences in this species [31].

In Experiment 2, we manipulated dissatisfaction during incubation, after all subject males had completed nest building. Nests were deemed complete once females began laying. Males built with breeder-standard coconut fibre, supplied in two 20-g piles on either side of the wall-mounted nestbox; fibre was replenished as needed during daily checks. Upon completion, males were pseudorandomly assigned to: (1) *Incubation-Satisfied* (Inc-Sat), allowed to incubate eggs, hatch chicks, and rear them to nutritional independence (∼ 35 days post-hatch; [26]), after which nests were removed and offspring separated at 40–45 days; or (2) *Incubation-Dissatisfied* (Inc-Diss), with all eggs removed during daily checks, preventing incubation. Assignment was structured in dyads: the first male to complete a nest went to Inc-Sat, the second to Inc-Diss. Within dyads, manipulation duration was matched. We weighed all removed nests, and this proxy for investment did not differ between treatments (mean grams *±* SE: Inc-Sat 39.71 *±* 12.49; Inc-Diss 39.61 *±* 13.89).

#### 2.2.2 Test: copy-if-dissatisfied?

##### Phase 1: preference test

One hour after light onset on the second test day, we gave each subject pair one pink and one orange string bundle (*n* = 25 pieces each; James Leaver Co., London, UK). To prevent nest building, bundles were tied to opposite cage sides, with side randomised across pairs. Four hours later, both were removed. From video, we quantified each male’s total time interacting—bill or feet—with each colour. If interaction with one or both colours was ≥ 60 seconds, the most-interacted-with colour was deemed preferred. If summed interaction time over the four hours was *<*60 seconds, the test was repeated the next day, and so on, until summed time across tests met our preference criterion (mean days ± SE to complete Phase 1: Mat-Sat 1.30 ± 0.21; Mat-Diss 1.00 ± 0.00; Inc-Sat 2.57 ± 0.43; Inc-Diss 2.53 ± 0.52).

##### Phase 2: pre-observation

After Phase 1, we moved a demonstrator pair into the unoccupied part of the test cage, allowing the demonstrator to begin nest building before being observed, as the opaque barrier remained in place. Phase 2 ensured that when observers later gained visual access: (1) demonstrators were actively building; and (2) all were at the same construction stage. We sham-tied 100 pieces of the subject male’s non-preferred colour midway along one cage wall and 50 pieces of his preferred colour on the opposite wall, with colour–side assignment randomised. The demonstrator could touch both colours but build only with the subject male’s non-preferred colour. The phase ended once all 100 non-preferred pieces were deposited in the nestbox (mean days ± SE to complete Phase 2: Mat-Sat 2.10 ± 0.57; Mat-Diss 1.56 ± 0.18; Inc-Sat 2.29 ± 0.16; Inc-Diss 2.40 ± 0.21).

##### Phase 3: observation

After Phase 2, a demonstrator pair received 50 pieces of the subject male’s nonpreferred material-type sham-tied to the cage wall. The 50 pieces of the preferred subject male’s material-type remained tied to the opposite cage wall. The 100 pieces of the subject male’s non-preferred material-type previously deposited in the nestbox remained in the nestbox. We then removed the opaque barrier between the cages, allowing the subject male visual access to the demonstrator male, until the demonstrator male had put all additional 50 pieces of the subject male’s non-preferred material-type into the nest (mean days ± SE to complete Phase 3: Mat-Sat: 1.28 ± 0.19; Mat-Diss: 1.06 ± 0.25; Inc-Sat: 1.25 ± 0.13; Inc-Diss: 1.20 ± 0.19.).

##### Phase 4: second-time nest building

After Phase 3, we replaced the opaque barrier and returned the demonstrator pair to colony housing. We then placed one pink and one orange string bundle (*n* = 25 pieces each) in the subject male’s cage, as in Phase 1. Phase 4 ended once he deposited all pieces of one colour in his nestbox. Material-choice decisions 1–25 were then scored from video. The demonstrated and non-demonstrated colours were considered the social and asocial materials, respectively.

### 2.3 Video scoring

All videos were scored blind to treatment using Boris 7.1.3 [41].

### 2.4 Statistical analyses

We analysed our data using `rstan` [42] and `rethinking` [43] in `R` [44]. After male drop-outs (because: *n* = 2 the female built; *n* = 2 the female died; *n* = 1 the female did not lay; *n* = 1 the video was corrupted), the final dataset included 47 subject males (*n* per treatment: Mat-Sat = 10; Mat-Diss = 8; Inc-Sat = 14; Inc-Diss = 15). Summary statistics above use this dataset.

#### 2.4.1 Behaviour models

To test how differential (dis)satisfaction affected males’ material choices, we compared (Model 1) first choices and (Model 2) all choices across treatments using hierarchical logistic regression. Both had individual-level random effects; Model 2 also modelled change over time via trial number and treatment-specific slopes, allowing variation across second-time nest building and between treatments.

#### 2.4.2 Cognition models

We built two EWA models: a baseline model without time-varying effects, and a monotonic model that includes time-varying effects. Fig. S1 shows parameter recovery tests validating each model *a priori* to data application.

##### EWA baseline model

This model consists of three components. The first component estimates males’ baseline nest-material attractions *A*_*i,j,t*=0_—that is, how preferable nest-material option *i* (pink or orange string) is to individual *j* during the initial material preference test (*t* =0):

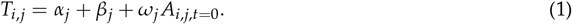

Here, *T* is the Poisson-distributed total number of touches to option *i* by individual *j*; *α*_*j*_ is an intercept term; *β*_*j*_ is a material exposure parameter, reflecting the individual-specific duration of Phase 1; and *ω*_*j*_ is a scaling parameter that weights individual *j*’s baseline attraction scores. Where touch data were missing (*n* = 19 males in Experiment 1, because this behaviour was not scored; we note that touch data in Experiment 2 were collected as part of a larger Master’s project), we did not exclude these individuals, as all parameters were estimated via partial pooling—borrowing information from the group (see Fig. S2).

The second component updates nest-material attractions during second-time nest building:

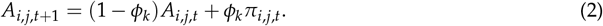

In words, attraction *A* to material *i* for individual *j* at time *t* + 1 is based on prior attraction *A*_*i,j,t*_ and the reward *π*_*i,j,t*_ from choosing that material at time *t*, weighted by *ϕ*_0 → 1_ for each treatment *k*. Larger values of *ϕ* indicate faster updating. Thus, *ϕ* represents *asocial information updating* of material preferences. We initialised *A*_*i,j,t*=0_ from the baseline estimates derived in Equation (1) such that *A*_social_ *< A*_asocial_. Following evidence that nest building activates reward circuits in zebra finches [45], we assigned a reward *π* =1 for choices and *π* =0 for non-choices.

The third component defines the probability of choosing material *i* by individual *j* at time *t* + 1 as a convex combination of asocial and social information:

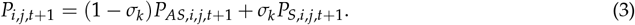

Here, *σ*_0 → 1_ controls the weighting of asocial versus social information. Larger values indicate greater consideration—but not necessarily adoption—of social information. Thus, *σ* represents *social information weighting*.

The probability of asocial influence is defined using a softmax function:

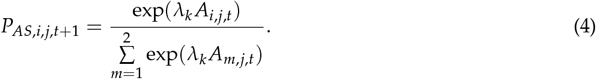

This transforms attraction scores into choice probabilities. The parameter *λ*_0 → ∞_ controls sensitivity to relative differences in attraction scores: higher values yield more deterministic choices, while *λ* =0 corresponds to random choice. Thus, *λ* represents *asocial information weighting* of material preferences.

The probability of social influence is based on a modified version of the frequency-dependent conformity function:

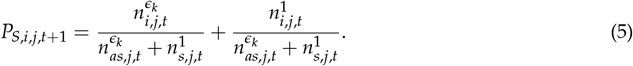

This formulation estimates the influence of deposited social material following demonstration via stimulus enhancement [46], adapted from [8]. Specifically, *n*_*as*_ and *n*_*s*_ refer to the cumulative number of asocial and social material pieces deposited by individual *j* in their nestbox. The parameter *ϵ*_0 → 1_ quantifies how reactive individuals are to changes in social-material abundance—conceptually akin to elasticity in economics:

- *ϵ >* 1: positively reactive (demand for social material increases disproportionately with its observed use)
- *ϵ <* 1: negatively reactive (demand for social material decreases disproportionately with its observed use)
- *ϵ* =1: unreactive (demand for social material is unaffected by changes in nestbox social-material abundance)

Thus, *ϵ* represents *social information updating*.

##### EWA monotonic model

The model structure remains as described above, with the key difference that each latent (a)social learning parameter—*ϕ, λ, ϵ*, and *σ*—varies over time. Following [14], we imposed a monotonic effects structure on each parameter *θ*, allowing the model to estimate per-trial step sizes—that is, the direction and magnitude of change across trials. This structure is given by:

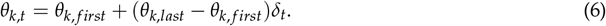

Here, *θ* _*f irst*_ and *θ*_*last*_ are treatment-level *k* endpoints, and *δ* is the cumulative sum of a 24-length simplex, representing the proportion of total change allocated up to trial *t*. This formulation ensures a monotonic— increasing or decreasing—trajectory over time, allowing the model to capture consistent learning trends as males progressed through second-time nest building.

Note, in both EWA models *λ* and *ϵ* were capped at 15 and 10, respectively, because larger values yield near-identical behavioural results and can introduce numerical instability.

#### 2.4.3 Evaluating EWA model performance

Predictive accuracy of EWA baseline and monotonic models was evaluated via approximate M-step-ahead predictions [47], testing time-series forecasts while preserving temporal dependencies. Analyses used `cmdstanr` [48] with custom R functions built on `loo` [49, 50] and `posterior` [51] packages (details in *Supplementary Material*).

#### 2.4.4 Forward simulations

We used agent-based forward simulations from the posterior of our EWA models to ask two questions (implementation details in *Supplementary Material*). While M-step-ahead forecasting tests predictive accuracy, forward simulations probe generative implications.

*Question 1: can we approximate our data?* It is possible that, in specifying the EWA models, we mischaracterised structural features of the decision-making process in ways that limit their ability to reproduce observed behaviour. Such mischaracterisations may not be evident from the estimated parameters alone. Instead, these parameters must be embedded in forward simulations to reveal potential mismatches between model assumptions and empirical patterns. To assess this, we matched the number of simulated agents to the sample size in each treatment and assigned each agent treatment-specific values for *ϕ, λ, ϵ*, and *σ*, sampled from the model’s random-effects multivariate normal distribution. This preserved the empirical correlation structure among parameters. We repeated this process 100 times to account for stochastic variability in simulated behaviour.

*Question 2: what would ‘choosers’ do in novel contexts?* Our EWA model estimates characterise a statistical, co-varying population of ‘choosers’. We used these estimates to explore how differential (dis)satisfaction might affect their use of asocial and social information in alternative choice contexts. Following the same simulation procedure as above, we simulated 10,000 ‘choosers’ (2,500 per treatment) and examined their behaviour in two hypothetical payoff structures: *social pays*, the social material pays a value of one, while the asocial material pays nothing; *social pays more*, the social material pays a value of two, and the asocial material pays a value of one. See the *Introduction* for the theoretical rationale behind this parameterisation.

## 3 Results

For all behavioural and computational modelling results, we report posterior means and 89% highest posterior density intervals (HPDI), representing the most probable parameter values given the model and data [10]. Logistic regression estimates reflect treatment-level effects incorporating individual variation via hierarchical structure but do not yield interpretable individual parameters. By contrast, EWA models explicitly estimate latent, individual-specific learning parameters (Fig. S3), which we summarise by treatment group.

### 3.1 Behaviour

Together, our behavioural results support a *copy-if-dissatisfied* strategy shaped by the relevance of social information to recent experience, with diminished evidence of the use of this strategy across material choices (Fig. 2).

**Figure 2.**
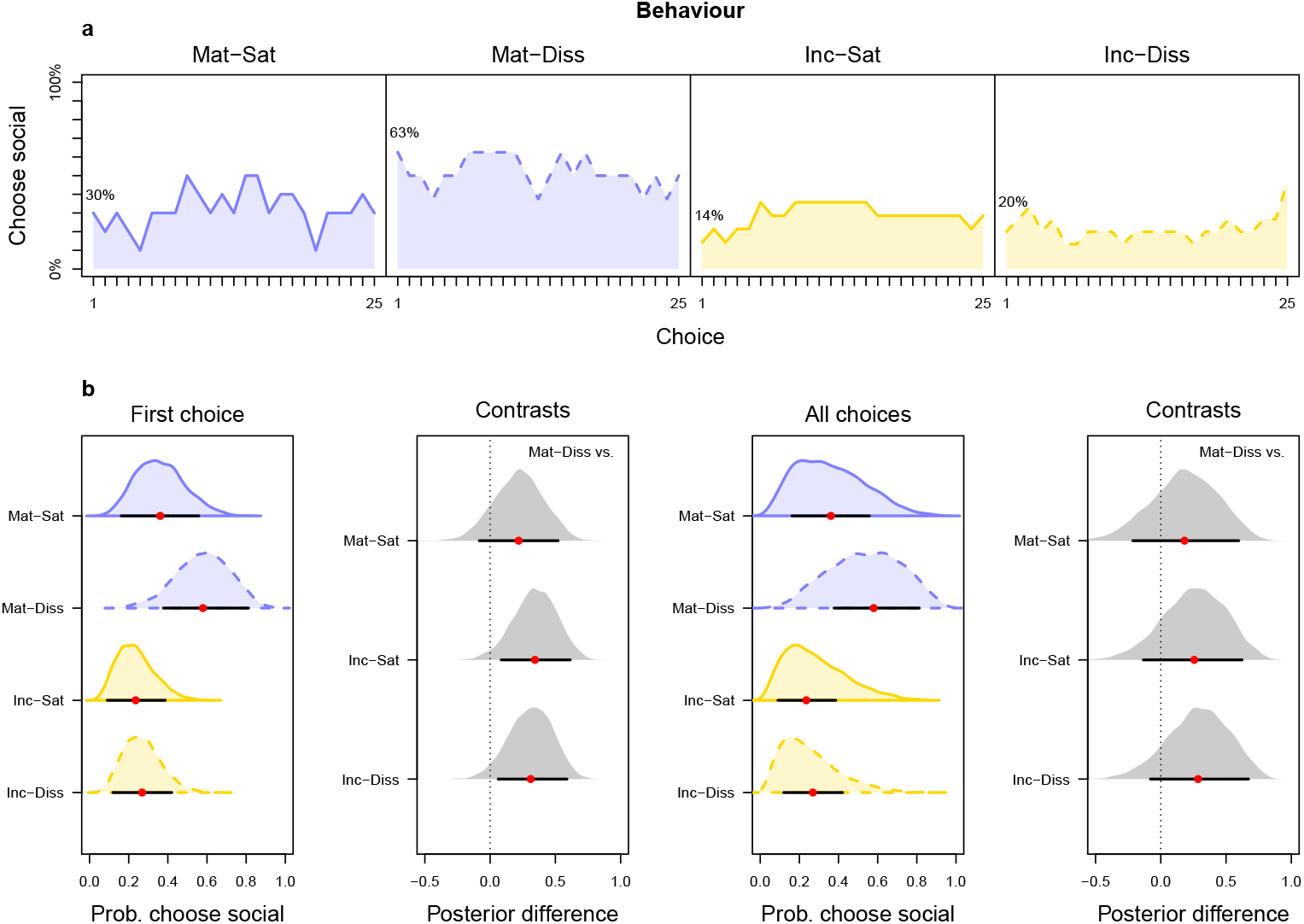
Behavioural results. **(a)** Mean percentage of males choosing social material over 25 choices per treatment, shown as lines with shaded areas for visual emphasis. **(b)** Density plots of posterior estimates (left to right): probability of choosing social material at first choice; contrasts between Mat-Diss and other treatments at first choice; overall probability across all choices; and corresponding contrasts. Red dots = posterior means; black lines = 89% HPDI.

#### 3.1.1 First choice

The decision to deposit social material first depended on a male’s prior experience: 63% of Mat-Diss males chose social material first—at least double the proportion observed in any other treatment (Mat-Sat: 30%; Inc-Sat: 14%; Inc-Diss: 20%; Fig. 2a). Posterior contrasts between Mat-Diss and the other treatments revealed an interactive effect of dissatisfaction and its type on males’ first material choice (Fig. 2b). Specifically, males in the Mat-Diss treatment—those given low-quality flexible string—were more likely to initially choose the social material compared to males in either (un)successful incubation treatment, as indicated by contrast intervals that excluded zero (Mat-Diss versus Inc-Sat: mean = 0.34, 89% HPDI: 0.09–0.62; Mat-Diss versus Inc-Diss: mean = 0.31, 89% HPDI: 0.04–0.57). A similar but weaker effect was observed in the comparison between males given flexible versus stiff string, where the contrast interval narrowly included zero (Mat-Diss versus Mat-Sat: mean = 0.21, 89% HPDI: − 0.08–0.52). Biologically, this pattern implies that it was specifically dissatisfaction with material that shifted initial choice toward the social material.

#### 3.1.2 All choices

Though less pronounced than at first choice, the influence of prior experience quality (satisfaction versus dissatisfaction) continued to interact with context (material versus incubation) to shape males’ material choices across the full nest-building sequence. Males in the Mat-Diss treatment continued to choose social material more frequently than males in the other treatments, although the magnitude of these differences was smaller (mean proportion of social deposits ± SE: Mat-Sat: 32% ± 10.62; Mat-Diss: 52% ± 13.52; Inc-Sat: 29% ± 11.23; Inc-Diss: 22% ± 8.76; Fig. 2a). Posterior contrasts between males in the Mat-Diss treatment and the other treatments supported this reduced effect (Fig. 2b): although between-treatment behavioural patterns persisted, they were strongest for the comparison with Inc-Diss (mean = 0.30, 89% HPDI: −0.07–0.67), weaker for Inc-Sat (mean = 0.26, 89% HPDI: −0.12–0.65), and weakest for Mat-Sat (mean = 0.18, 89% HPDI: −0.21–0.58).

### 3.2 Cognition

Together, our cognitive modelling results reveal the dynamic nature of avian nest building: males’ material choices were shaped by plastic learning mechanisms that balanced asocial and social cues in response to recent outcomes. All parameter estimates are reported in Table S1; we summarise the key results below.

#### 3.2.1 Asocial information updating (*ϕ*): how rapidly did males update material preferences?

The EWA baseline model estimates for *ϕ* were moderate across treatments (Fig. 3, row one, left panels), indicating reasonably fast material preference updating by males. In the EWA monotonic model (Fig. 3, row one, right panels), *ϕ* remained fairly stable across choices, with a slight decline among most males, except those in the Inc-Diss treatment, who showed a small increase. These time-structured results suggest little overall change in how rapidly males updated material preferences over nest building.

**Figure 3.**
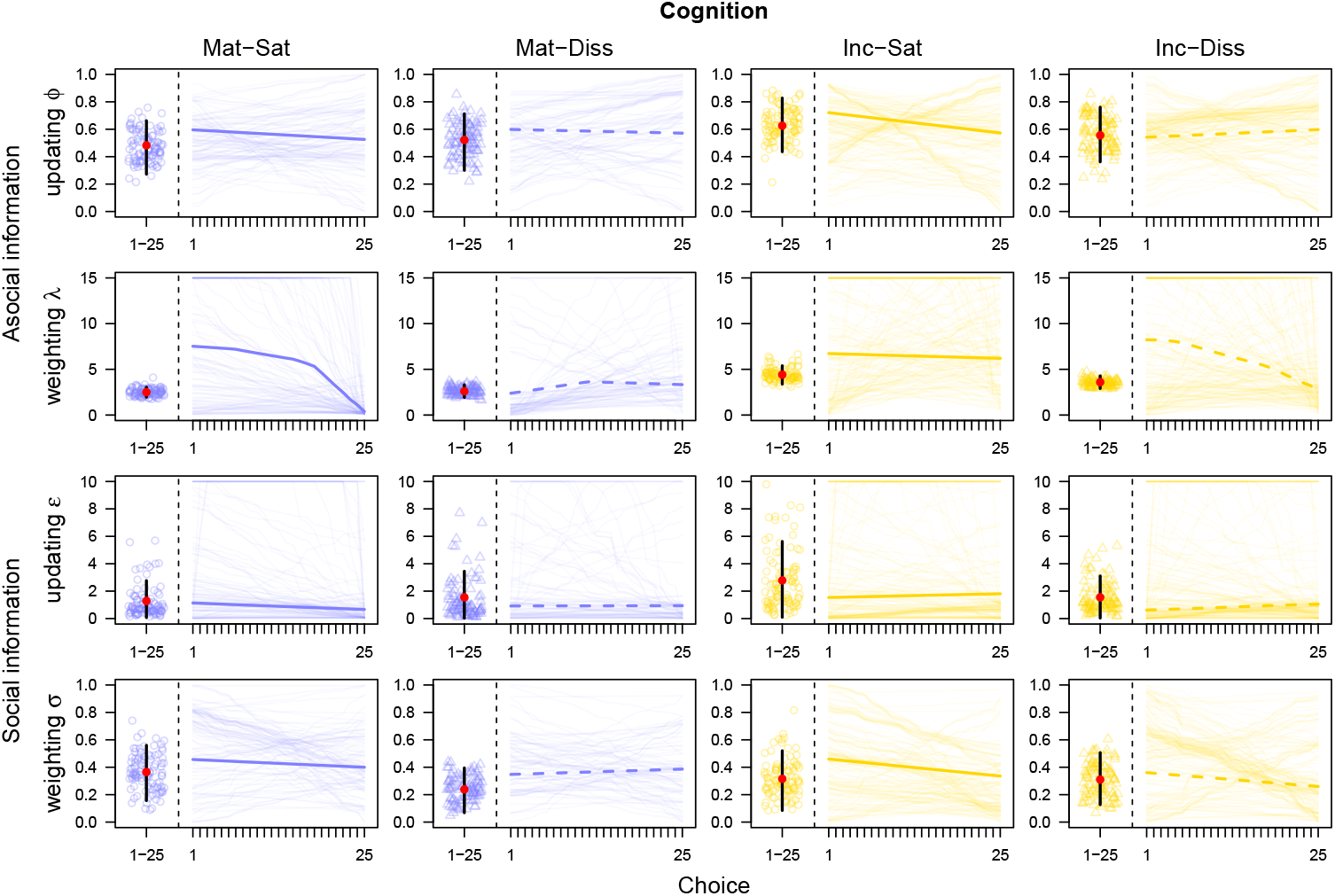
Cognition results. Latent (a)social learning parameters *ϕ, λ, ϵ*, and *σ* by treatment and EWA model. Left panels: baseline model estimates with posterior means (red dots), 89% HPDI (black lines), and 100 posterior draws (open circles/triangles). Right panels: monotonic model estimates with posterior means (solid/dashed lines) and 10 posterior learning trajectories per male (thin lines). Fig. S3 shows these results at individual-level resolution.

#### 3.2.2 Asocial information weighting (*λ*): how strongly did males exhibit material preferences?

The EWA baseline model estimates for *λ* were highest in the incubation treatments and moderate in the material treatments, indicating stronger material preference expression following reproductive experience (Fig. 3, row two, left panels). In the EWA monotonic model (Fig. 3, row two, right panels), Mat-Sat males showed a sharp decline in *λ* over time, suggesting a shift toward more exploratory decisions, as reflected by their increased material-choice stochasticity (Fig. 2a). For Mat-Diss males, *λ* began at a moderate level and increased moderately over time, indicating increasingly stable, preference-based decisions; in this case, directed toward the social material (Fig. 2a). Both incubation treatments maintained high *λ* values, consistent with their sustained majority-use of the asocial material (Fig. 2a). However, Inc-Diss males exhibited a gradual decline in *λ*, paralleling their late-trial small spike in social material choice (Fig. 2a).

#### 3.2.3 Social information updating (*ϵ*): how elastically reactive were males to deposited social material?

The EWA baseline model estimates for *ϵ* revealed weak positive (*>*1) social reactivity across males (Fig. 3, row three, left panels), with Inc-Sat individuals showing the strongest estimates, consistent with their early up-tick in choice for social material that then plateaued (Fig. 2a). These results suggests a broad but subtle tendency for demand for social material to increases disproportionately with its observed use. The EWA monotonic model showed that this stimulus enhancement effect remained relatively stable throughout nest building (Fig. 3, row three, right panels), with *ϵ* showing only slight upward or downward trends—hovering just above, below, or crossing 1 depending on treatment.

#### 3.2.4 Social information weighting (*σ*): how much did males consider asocial versus social information?

The EWA baseline model estimates for *σ* were moderate across treatments (Fig. 3, row four, left panels), suggesting predominant consideration of asocial information when choosing material. In the monotonic model (Fig. 3, row four, right panels), *σ* remained relatively stable in the material treatments but declined somewhat over time in both incubation treatments. Overall, these time-structured results show a sustained, though modest, consideration of social information despite an absence of strong conformity patterns (Fig. 2a)

### 3.3 Performance

Using an approximation technique to forecast EWA model choices, both baseline and monotonic models performed reliably (Fig. S4, top row), with a slight predictive advantage for the baseline model (Fig. S4, bottom row).

### 3.4 Simulations

#### 3.4.1 Approximation

Together, our forward simulations—matched to treatment sample sizes—show that the EWA baseline model approximated our nest-building data more effectively than the EWA monotonic model, though the difference was modest (Fig. S5). In general, simulated ‘choosers’ tended to use more of the social material than their real-world counterparts, except in the Mat-Diss treatment where mapping was fairly close. This overestimation likely reflects two structural constraints. First, the reward structure assumes that each choice is equally reinforcing, potentially exaggerating the attractiveness of the social material. Second, in the monotonic model, this reinforcing effect is combined with the imposed cumulative change in parameters over time. In both cases, these constraints may steepen learning trajectories, causing the simulations to overshoot observed social material use. Accordingly, we rely on the more behaviourally-aligned EWA baseline model when probing generalisation beyond our study, recognising that the true value of material choice is likely more individual-specific and nuanced.

#### 3.4.2 Generalisation

Building on the above simulations, we extended our forward simulations to explore two plausible realworld material-choice contexts that deliberately varied in their reward structure (Fig. 4). Here, rather than assuming equal reinforcement, we explicitly tested how differential rewards for social versus asocial material might shape behaviour. Most notably, simulated ‘choosers’ were more likely to select the social material in the *social pays* context—where only the social material yielded a reward—than in the *social pays more* context, where the social material paid twice as much as the asocial option. Overall, (dis)satisfaction had little effect on choices by ‘choosers’. However, recent nest-building experience— rather than incubation experience—tended to reliably shift choice toward the social material, suggesting that sensitivity to the relevance of social information may generalise beyond the original experimental context.

**Figure 4.**
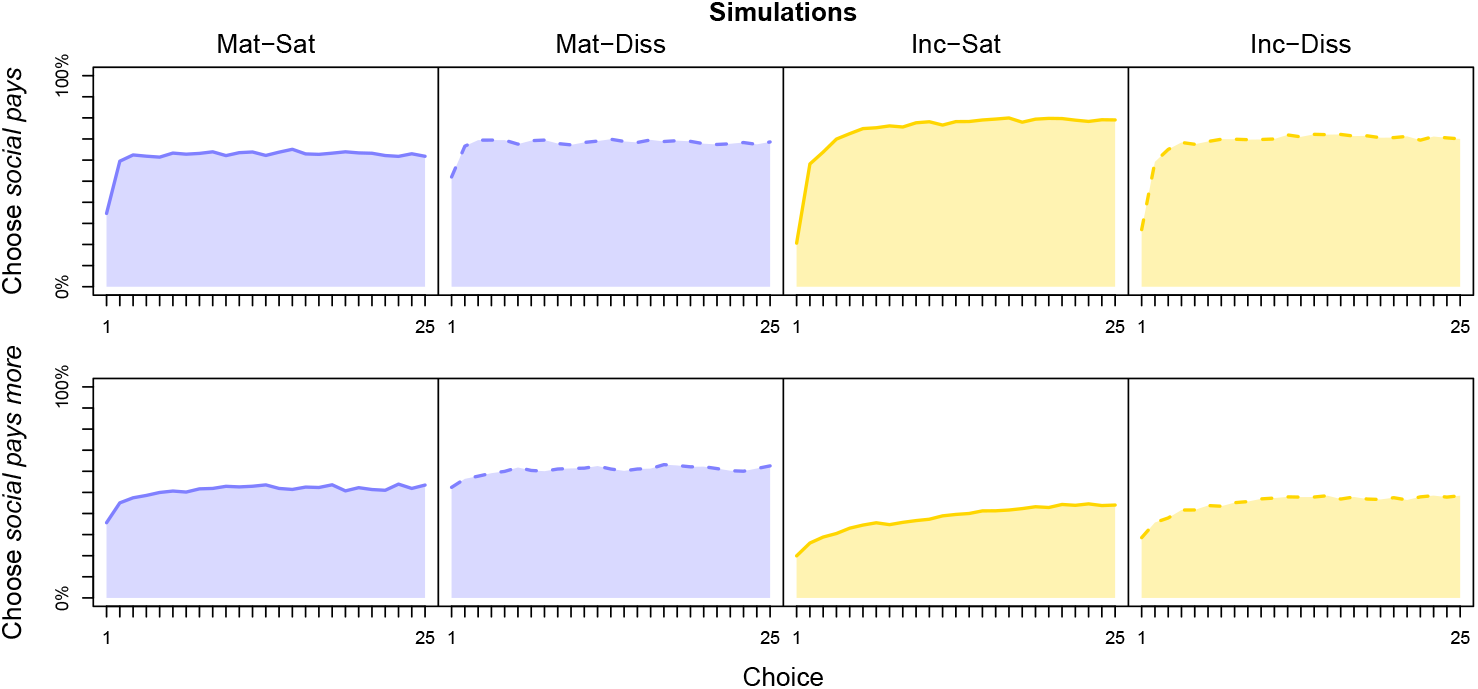
Simulation results. Mean percentage of ‘choosers’ (*n* = 2,500 per treatment) using social material across 25 choices under varying reward contexts, shown as lines with shaded areas for visual emphasis. Top row: social material paid one, asocial paid nothing (*social pays*). Bottom row: social material paid two, asocial paid one (*social pays more*).

## 4 Discussion

Using a combination of behavioural experiments, Bayesian cognitive modelling, and forward simulations, we set out to examine strategic social learning in nest-building zebra finches—specifically testing the *copy-if-dissatisfied* strategy, and modelling the underlying cognitive mechanisms plus their potential generalisability. Below, we discuss our behavioural results, cognitive model findings, and exploratory simulation outcomes.

### 4.1 Nest-building zebra finches *copy-if-dissatisfied* with material experience

After observing conspecific nest building, zebra finches chose more of the demonstrated social material— particularly for their first material choice—if they had previously experienced low-quality material during nest building. Notably, neither high-quality material nest-building experience nor reproductive outcomes—whether successful or unsuccessful in incubation—led to as much social material use as that observed in males with poor nest-building material-experience. Together, these behavioural results suggest nest-building zebra finches are more likely to use social information about ‘what to choose’, especially initially, when their own material-use experience has been dissatisfactory, consistent with a *copy-if-dissatisfied* strategy.

The influence of individual dissatisfaction on social information use has previously been demonstrated experimentally in several taxa, including bats [52], beetles [53], bumblebees [54], and rats [55]. More broadly, the use of strategic social learning rules to filter information and optimise learning opportunities is both theoretically and empirically supported—examples include biasing social learning toward kin or adults [56, 57], or monitoring the frequency or reward-payoffs of others’ choices [13, 17]. We extend this general literature by showing not only that dissatisfied animals use social information strategically, but also that this strategising appears adaptive—that is, fine-tuned to expected utility.

Here, *utility* refers to the potential benefit of copying conspecific-presented nest material. Given that zebra finches associate nest-building efficacy with material properties—such as rigidity, which may influence reproductive success [31, 32]—our finding that males with prior experience using ‘bad’ flexible string tended to choose more of the demonstrated social material is consistent with this predicted association. In contrast and against our prediction, males whose clutches were removed used less social material, suggesting they may have attributed their reproductive failure to other causes, such as nest predation triggered by the observed clutch removal (birds can learn such cause-effect associations: [21, 58]). If so, different types of social information—such as cues about where to nest—may have been more relevant to them, a hypothesis open to future experimental testing. The relatively low use of social material by satisfied males likely reflects the absence of an adverse first-time building or incubation experience, consistent with a simple win-stay heuristic. Future research could investigate whether this heuristic breaks down when characteristics of demonstrators, such as relatedness or age [1], are experimentally manipulated.

Nest-building zebra finches, then, did not apply a blanket *copy-if-dissatisfied* strategy, nor did they copy exclusively when they did. We observed individual variation in social material use over time, with the highest observed use being only 63% by Mat-Diss males on their first material choice. This relatively modest initial level of copying may reflect the absence of any functional—and thus reinforcing— difference between the asocial and social material [33]. Importantly, the individual-level and temporal variation we observed underscores the dynamic nature of asocial and social information use—a process our cognitive modelling aimed to capture.

### 4.2 Nest-building zebra finches dynamically inform their material choice

We developed dynamic EWA learning models [8, 36] to estimate how latent asocial and social learning parameters shaped nest-building zebra finches’ material choices, including one version that allowed parameters to change over time. However, the time-structured model was marginally less successful at predicting and approximating the real behavioural data (see Sections 3.3 and 3.4.1), so we focus here on results from the more robust baseline model.

All zebra finches exhibited moderately high *asocial information updating* (*ϕ*)—they readily revised their material preferences based on recent material choices. This result aligns with one of the only other studies, to our knowledge, that estimated *ϕ* in birds within a social learning context, which similarly found high responsiveness to individual choice experience, although this declined with age [13]. While our study was not designed to test age effects, the fact that zebra finches learn nest building both asocially and socially [21, 22, 33, 59] suggests it is plausible that resistance to new nest-related information increases with age. This possibility merits further investigation, particularly since recent EWA simulations show *ϕ* can influence the speed at which socially acquired behaviours are produced [60], implying that younger birds may conform more quickly than older individuals to local architectural traditions [23, 25, 37].

Estimates for *asocial information weighting* (*λ*) were high across all zebra finches and even higher in the incubation treatments than in the material treatments. This implies that males generally exhibited strong material preferences and switched infrequently between materials. In the computational reinforcement learning literature[61, 62], such choice patterns are typically interpreted as indicators of high reward sensitivity or low exploration—reflecting risk-sensitive or deterministic behaviour [11, 14, 60]. Although our EWA model rewarded males equally for both materials (see *Statistical analyses*), zebra finches received no explicit material-choice rewards during the study. However, nest building has been shown to activate reward circuits in zebra finches [45], suggesting dopaminergic mechanisms may underlie our males’ apparently conservative material preferences—in other words, males may have largely stuck with the material they ‘liked’.

All zebra finches exhibited modest (*>*1) *social information updating* (*ϵ*): with each social material deposit, they became slightly more likely to choose it, indicating a tendency to demand more of this socially-enhanced stimulus rather than ignoring or reducing interest. This stimulus enhancement effect could be estimated due to the adaptability of the EWA modelling framework, which accommodates a wide range of mathematically defined processes—including age- or payoff-based biases [8, 14, 16] —and can be tailored to contexts like ours, where social demonstration occurs outside of group-level conformity dynamics. More broadly, computational modelling encourages researchers to formalise hypotheses about learning processes, supporting more precise and testable accounts of behaviour than those offered by catch-all terms; for example, ‘behavioural flexibility’ [11, 63].

Finally, when comparing *social information weighting* (*σ*) to the asocial parameters (*ϕ* and *λ*), zebra finches overall placed greater weight on asocial information, though they did consider both types of cues. That the EWA model captured these nuanced influences represents a methodological advance over previous work on zebra finches, where—lacking evidence of strong conformity—social influence was often inferred from initial material touches alone [33, 59, 64]. Our generative forward simulations, discussed below, suggest in contexts where social material confers a clear payoff advantage, consideration of social information could outweigh that of asocial cues more strongly than in the present study.

### 4.3 Beyond nest-building zebra finches

In simulations exploring how differential (dis)satisfaction influenced choice behaviour in new contexts where rewards varied by material, we found that this prior experience had minimal impact; instead, choices were primarily driven by payoffs.

Perhaps the most striking result emerged in the *social pays* context, where only the social material yielded a reward: here, ‘choosers’ selected the most social material. This pattern highlights how payoff-sensitive *asocial information updating* (*ϕ*) can steer behaviour away from costlier options—since choice behaviour feeds back to shape material preferences (see Equations 2 and 4)—and suggests population-level (cultural) differences in material use (e.g., [24, 37, 38, 39]) can emerge even without explicit strategic social learning rules. Supporting this, experimental data show combining asocial payoff sensitivity with conformity bias can produce adaptive foraging traditions in wild birds without the need for strategising based on others’ payoffs [13].

Notably, in the *social pays more* context—where the social material offered twice the reward of the asocial one—’choosers’ following (un)successful incubation used less social material than in the *social pays* context. This counterintuitive result suggests simply increasing the relative payoff of a social material-option may not be sufficient to drive stronger use of it. In humans, cumulative improvements in behaviour are thought to result from a combination of mechanisms—including copying, innovation, exploration, and biases such as success- or prestige-bias—as well as environmental factors like spatial variability [14, 65, 66]. A similarly multi-faceted process is thus likely required to scaffold supposed cumulative improvements in material-use behaviour in animals (e.g., [39]). For example, wild birds learning a high-reward, multi-step foraging task rely on social learning for the lower-reward components, then asocially chain those steps together through practice [67]. Likewise, wild birds temporarily housed in captivity attend more closely to social information about reward differences when they have recently come from a different habitat [16].

Our simulation-derived data should, however, be interpreted with caution, given the limited sample—both in size and taxonomic scope—and the reductionist structure of the computational setup. Future work should improve on these limitations, maybe beginning with better individual-level approximations of material-choice evaluations.

### 4.4 Conclusions

Once thought entirely innate, avian nest building is now recognised as shaped by individual learning, with growing evidence for social influences and, more recently, architectural culture [21, 22, 23, 24, 25, 59]. We show that nest-building zebra finches follow a *copy-if-dissatisfied* strategic social learning rule, but only when social information aligns with recent experience—suggesting recognition of its utility. We identified the latent learning mechanisms driving these decisions—two asocial and two social parameters—thereby confirming the cognitive basis of avian nest building. We validated these mechanisms with generative simulations, which showed the baseline model generalised slightly better than the time-varying alternative, consistent with predictive performance checks. This underscores the value of simulation-based falsification for identifying cognitive models that best capture real-world decision-making [9]. Extending the more robust, behaviourally aligned model to simulated ‘choosers’ in novel reward contexts produced preliminary predictions about drivers of material-use variation, offering a foundation for comparative work across taxa and contexts. Our open-access analytical approach and insights, alongside recognition of cognitive and phenotypic parallels between nest building and tool use [21, 23, 25, 68, 69, 70], aim to stimulate research into material-use learning dynamics, particularly from a computational framework [15].

## Acknowledgements

We are grateful to two anonymous reviewers for their thoughtful feedback, which improved our manuscript. A.J.B. extends heartfelt thanks to (Prof./Pops) Stephen Breen for productive conversations on structuring Equation (5). A.J.B. further thanks (Dr.) Dominik Deffner for sharing code on GitHub, which was an invaluable resource.

## Supplementary material for the manuscript

### Further details on housing and husbandry

Each demonstrator pair were, at the start of the current study, experienced builders and already paired together: because the male of the demonstrator pair had previously provided social artefact nests in an unrelated study. None of the demonstrator pairs were the parents or the siblings of the subject pairs. Likewise, no subject pair consisted of siblings.

When not participating in the current study, all subject and demonstrator male-female zebra finch pairs were housed in same-sex free-flight colony housing; more specifically, either in colony rooms (Experiment 1: males, 318 × 312 × 230 cm; females, 438 × 251 × 230 cm) or in colony cages (Experiment 2: males and females, 165 × 66 × 184 cm).

During their respective experiment, subject male-female pairs were first housed in training cages during their experiment-specific dissatisfaction manipulation (Experiment 1: 75 × 75 × 40 cm; Experiment 2: 50 × 50 × 100 cm). After this, they were housed in testing cages (Experiment 1 and 2: 50 × 50 × 100 cm). Demonstrator male-female pairs, by contrast, were only housed in testing cages. Training cages were placed together in one, or between two, experimental room(s) (Experiment 1 and 2, respectively). In either case, subject male-female pairs could only hear and not see one another due to opaque between-training-cage barriers. Testing cages were placed in a pair in one, or in three different, experimental room(s) (Experiment 1 and 2, respectively), roughly 10 cm apart and back-to-back, with a removable opaque barrier allowing for control of visual access between sets of testing cages. Depending on the experiment, training and testing cages were either fitted with, respectively, one and three cameras (Experiment 1; Spy Camera CCTV, Bristol, United Kingdom), or, each with three cameras (Experiment 2; mini BNC cameras, OSY CAMS), to habituate or record subject male’s behaviour on a corresponding laptop computer. All training and testing cages each contained six perches, two food and two water bowls, and a wooden open-faced nestbox (both experiments: 12.5 × 12 × 12 cm), which was hung midway on the back cage panel.

Irrespective of housing, all subject and demonstrator male-female zebra finch pairs had *ad libitum* access to seed, vitamin-supplemented water, oystershell grit, cuttlefish bone, and vitamin block; plus fresh spinach and/or spray millet three times per week. The zebra finches were kept at approximately 19–24°C, on a 14:10 light:dark hour cycle, with humidity levels between 30–50%. Finally, Experiment 2 subject male-female pairs (plus their offspring) received fresh egg mix, daily.

### Further details on the predictive power checks

We assessed predictive power using approximate M-step-ahead predictions, specifically Pareto-Smoothed Importance Sampling Leave-Future-Out Cross-Validation (PSIS-LFO-CV), as outlined by Bü rkner, Gabry, and Vehtari in the the following vignette: https://mc-stan.org/loo/articles/loo2-lfo.html. This procedure offers two key advantages: (1) it provides a time-sensitive evaluation of model fit that respects the temporal structure of learning, and (2) it enables direct comparison of time-varying models.

In brief, PSIS-LFO-CV extends traditional cross-validation methods to time series by holding out *future* observations rather than individual points. This respects temporal dependencies by training the model on a moving window (e.g., trials 1 to *t*) and assessing its ability to predict data *M* steps ahead (e.g., *t* + 1 when *M* = 1). While these future values are known, the model is asked to predict them as if they were not—simulating true out-of-sample prediction. Note that the final observations in a time series cannot be evaluated in this way, as there is no future data for comparison.

Crucially, PSIS-LFO-CV allows us to approximate these predictions without re-fitting the model at every step. Instead, we use existing posterior samples and apply Pareto smoothing to stabilise importance weights, especially in cases with high variance. When the Pareto-*k* diagnostic exceeds 0.7, this suggests that the approximation may be unreliable, and full re-fitting may be warranted. It is possible to track these diagnostics and to report the proportion of windows requiring re-fitting.

In addition to individual model checks, PSIS-LFO-CV enables direct model comparisons via differences in expected log predictive density (ΔELPD), which quantify how much better one model predicts future data compared to another. Specifically, ΔELPD is computed as Model 1 ELPD minus Model 2 ELPD; negative and positive values thus indicate an advantage for Model 1 or Model 2, respectively.

For each model, we implemented a moving-window procedure that began after an initial training window *L* = 5 trials per subject male—an intentionally conservative window designed to stress-test the EWA models’ ability to predict from limited data. At each step *t*, we assessed predictive performance on the next trial (*t* + 1) using 1-step-ahead log predictive density estimates (i.e., with forecasting horizon *M* = 1). Log-likelihoods were computed from posterior samples obtained by fitting the model to trials 1 through *t*. The final trial (*t* = 25) was excluded from analysis, as it lacked a subsequent observation against which predictions could be compared. When diagnostics indicated poor approximation (as given by Pareto-*k >* 0.7), the model was automatically re-fit on the training window to ensure stable and reliable estimates.

Here, we recorded the Pareto-*k* values and the number of re-fits, providing diagnostics of predictive reliability and computational stability. To compare models, we computed the ELPD for each individual and trial based on the rolling PSIS-LFO-CV procedure described above, and then calculated ΔELPD (monotonic ELPD − baseline ELPD) to quantify relative predictive performance. Results are summarised in Fig. S4. All forecasting code is openly available at our GitHub repository: https://github.com/alexisbreen/Dynamic-nest-building/tree/main.

### Further details on the forward simulation technique

To probe the generative adequacy of our model and test its predictions under novel contexts, we implemented forward simulations using posterior draws from our hierarchical EWA models. Below we detail how these simulations were constructed.

#### Simulation structure

For each simulation run, we generated a set of virtual agents (‘choosers’) matched in number to each treatment’s empirical sample size (for data-approximation simulations; hereafter S1) or scaled to 2,500 per treatment (for generalisation simulations; hereafter S2). Each ‘chooser’ was assigned a set of fixed latent learning parameters: in S1, *ϕ, λ, ϵ*, and *σ*; in S2, *ϕ*_first_, *ϕ*_last_, *λ*_first_, *λ*_last_, *ϵ*_first_, *ϵ*_last_, *σ*_first_, *σ*_last_, and *δ*. These values were independently drawn from the posterior distribution of the treatment-specific individual-level parameters *θ*_*k*_, estimated across 4,000 iterations by our EWA models.

For example, a ‘chooser’ in S1 from the Mat-Sat treatment (*k* = 1) was randomly assigned a draw *d* from iteration three, resulting in the assignment of *ϕ*_*k*=1,*d*=3_, *λ*_*k*=1,*d*=3_, *ϵ*_*k*=1,*d*=3_, and *σ*_*k*=1,*d*=3_ to govern their behaviour. In this way, we preserved treatment-level variation and the posterior correlation structure among parameters within treatments in both S1 and S2.

#### Choice updating process

Each ‘chooser’ began with initial attraction values *A* for the two material types *i* ∈ {social, asocial }, derived from the empirical estimates of Equation 2 at the first trial. Continuing the example above, the S1 Mat-Sat ‘chooser’ would receive:

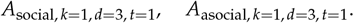

We drew from the estimated attraction scores at *t* = 1 rather than from the baseline estimates (i.e., Equation 1), as these values more accurately reflect true post-demonstration attractions due to the recursive structure of the estimation process. These initial attractions were then updated across 25 simulated trials using Equation 2, with learning dynamics governed by the assigned *ϕ, λ, ϵ*, and *σ* values (or their timevarying counterparts in S2). In other words, material preferences evolved across trials due to cumulative updating, ensuring that choosers were not statically assigned to prefer one material throughout.

#### Payoff structures and reward assignment

We manipulated the reward term *π* as follows:

- In S1, *π* = 1 for a chosen option and *π* = 0 for the non-chosen option, regardless of whether it was the asocial or social material. This mimics the reinforcement potential of nest building as an intrinsically rewarding behaviour.
- In the S2 *social pays* context, we set *π* = 1 for choosing the social material and *π* = 0 for the asocial material.
- In the S2 *social pays more* context, *π* = 2 for choosing the social material and *π* = 1 for the asocial material.

In all cases, these *π* values directly influenced subsequent attraction updates via Equation 2 and thereby shaped evolving material preferences and choice probabilities over time.

### Summary

To account for stochasticity we ran 100 replicate simulations matched to study sample sizes for S1, and 10,000 simulations (2,500 choosers per treatment) in S2. In each simulation, all ‘choosers’ progressed through 25 simulated material choices, dynamically updating their preferences using the same EWA mathematical structure as in the empirical model, with choice payoffs predetermined by simulation and context. Latent learning parameters were fixed per ‘chooser’, drawn at random from the posterior distribution, and preserved the empirical correlation structure within treatments at the individual-level. All simulation code is openly available at our GitHub repository: https://github.com/alexisbreen/Dynamic-nest-building/tree/main.

**Table S1.** Posterior means and 89% lower (Lo) and higher (Hi) posterior density intervals for each latent learning parameter, by treatment and model. 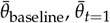, and 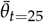 denote the average of individual-specific parameter estimates from the EWA baseline model, and from material choice 1 and 25 in the EWA monotonic model, respectively.

| Treatment | Parameter | $\bar{\theta}_{\text{baseline}}$ | Lo | Hi | $\bar{\theta}_{t=1}$ | Lo | Hi | $\bar{\theta}_{t=25}$ | Lo | Hi |
| --- | --- | --- | --- | --- | --- | --- | --- | --- | --- | --- |
| Mat-Sat | $\phi$ | 0.49 | 0.29 | 0.68 | 0.60 | 0.56 | 0.66 | 0.53 | 0.49 | 0.56 |
| Mat-Diss | $\phi$ | 0.53 | 0.32 | 0.74 | 0.60 | 0.56 | 0.64 | 0.57 | 0.50 | 0.63 |
| Inc-Sat | $\phi$ | 0.63 | 0.43 | 0.82 | 0.72 | 0.70 | 0.76 | 0.57 | 0.53 | 0.58 |
| Inc-Diss | $\phi$ | 0.55 | 0.36 | 0.74 | 0.54 | 0.49 | 0.58 | 0.60 | 0.54 | 0.69 |
| Mat-Sat | $\lambda$ | 2.48 | 1.94 | 3.00 | 7.54 | 0.36 | 15.00 | 0.37 | 0.23 | 0.70 |
| Mat-Diss | $\lambda$ | 2.62 | 1.96 | 3.35 | 2.35 | 0.18 | 15.00 | 3.36 | 0.85 | 5.90 |
| Inc-Sat | $\lambda$ | 4.40 | 3.44 | 5.45 | 6.62 | 0.36 | 10.90 | 6.24 | 5.45 | 9.81 |
| Inc-Diss | $\lambda$ | 3.59 | 2.94 | 4.27 | 8.21 | 0.44 | 15.00 | 2.83 | 0.84 | 5.15 |
| Mat-Sat | $\epsilon$ | 1.26 | 0.08 | 2.72 | 1.15 | 0.64 | 2.37 | 0.65 | 0.09 | 2.80 |
| Mat-Diss | $\epsilon$ | 1.50 | 0.06 | 3.48 | 0.97 | 0.38 | 1.74 | 0.83 | 0.12 | 2.90 |
| Inc-Sat | $\epsilon$ | 2.84 | 0.11 | 5.77 | 1.55 | 1.36 | 2.46 | 1.83 | 0.19 | 4.50 |
| Inc-Diss | $\epsilon$ | 1.53 | 0.09 | 3.06 | 0.64 | 0.30 | 0.80 | 1.06 | 0.15 | 1.57 |
| Mat-Sat | $\sigma$ | 0.36 | 0.15 | 0.56 | 0.45 | 0.33 | 0.49 | 0.40 | 0.34 | 0.47 |
| Mat-Diss | $\sigma$ | 0.23 | 0.06 | 0.38 | 0.34 | 0.28 | 0.45 | 0.39 | 0.32 | 0.45 |
| Inc-Sat | $\sigma$ | 0.31 | 0.09 | 0.52 | 0.46 | 0.40 | 0.52 | 0.33 | 0.31 | 0.37 |
| Inc-Diss | $\sigma$ | 0.31 | 0.11 | 0.48 | 0.36 | 0.28 | 0.40 | 0.26 | 0.19 | 0.30 |

**Figure S1.**
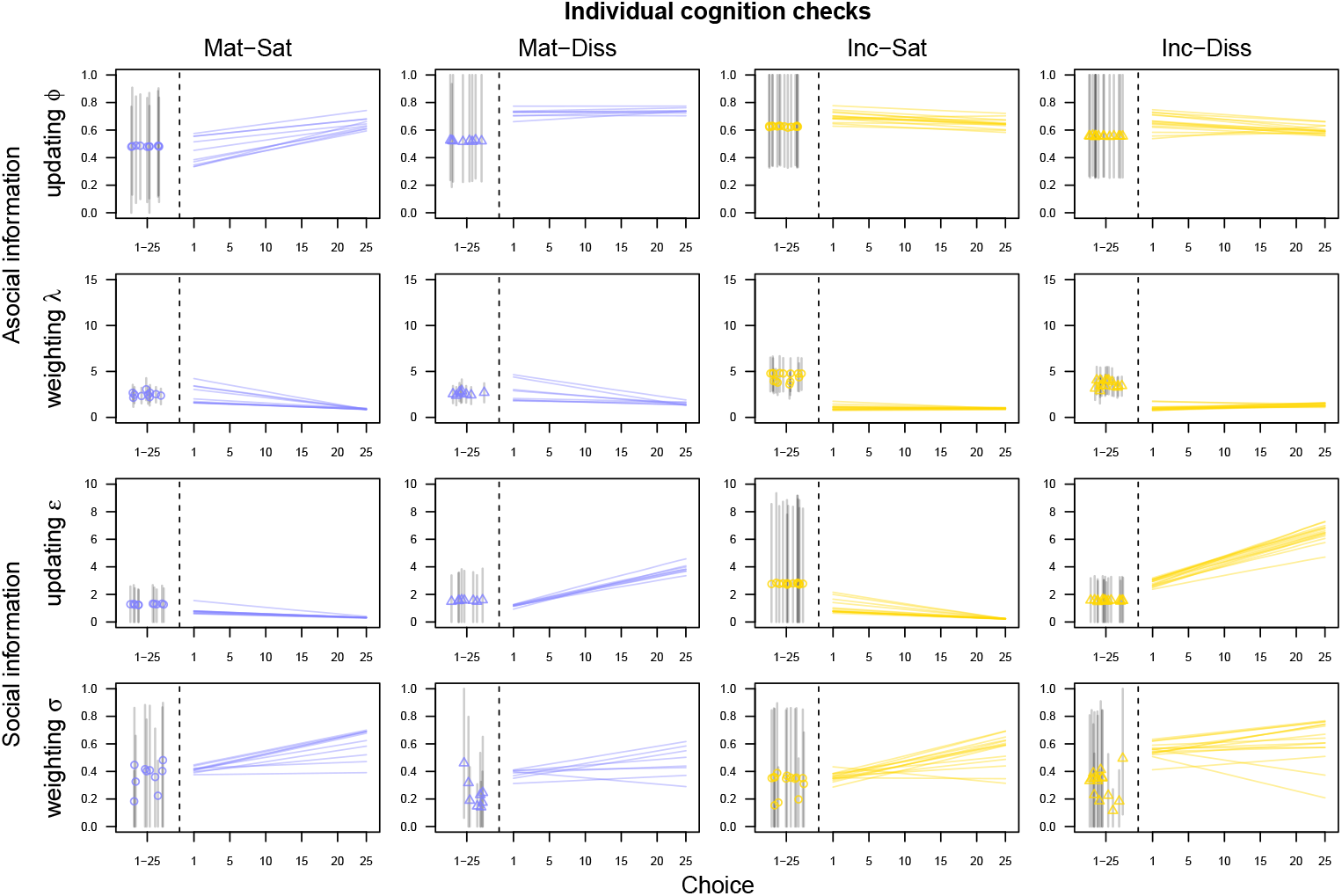
EWA model validation. Prior to fitting our EWA baseline and monotonic models to the empirical data, we validated both models via standard parameter recovery tests using simulated data matched to our treatment sample sizes. This figure shows one such test. For the baseline EWA model, parameters were specified as: *ϕ* = 0.5, *λ* = 3, *ϵ* = 1, and *σ* = 0.5; with both between and within treatment, and between experiment variation. For the monotonic EWA model, all parameters were identical except for *ϵ*, the elasticity of demand for social material, which was designed to vary over time. Specifically, to model a differential (dis)satisfaction effect, we simulated distinct behaviours for satisfied and dissatisfied ‘choosers’: Mat-Sat and Inc-Sat ‘choosers’ decreased their demand by approximately 80%, while Mat-Diss and Inc-Diss ‘choosers’ increased their demand by approximately 500%. Both models recovered these specification dynamics within a reasonable range. Left panels display EWA baseline model estimates: open circles or triangles represent posterior means for each ‘chooser’, with black lines indicating 89% highest posterior density intervals. Right panels display EWA monotonic model estimates: each line represents the posterior mean trajectory across 25 choices for a single ‘chooser’.

**Figure S2.**
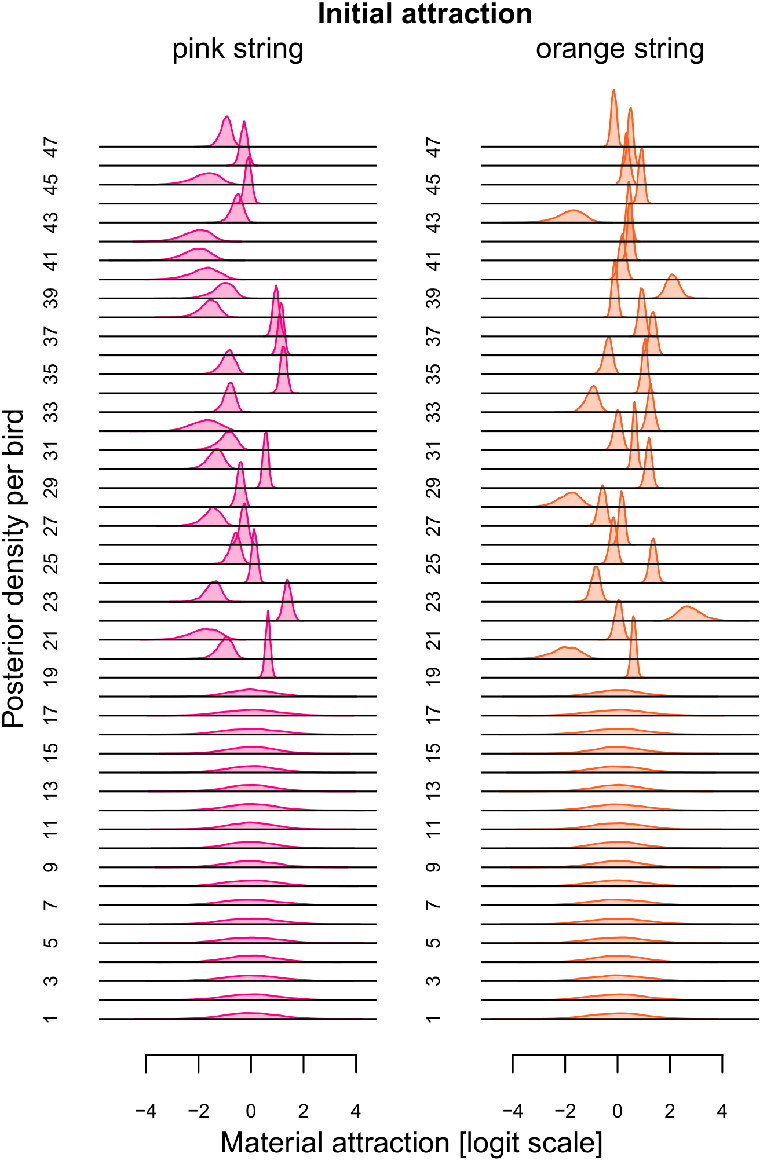
Initial material attractions. Density plots showing posterior distributions of initial pink and orange string attractions for subject males (see Equation 1 in the main text). Note that the wider distributions for Birds 1–18 reflect the absence of touch data from Experiment 1. Estimation for these individuals remains possible through partial pooling, which borrows strength from the group-level structure.

**Figure S3.**
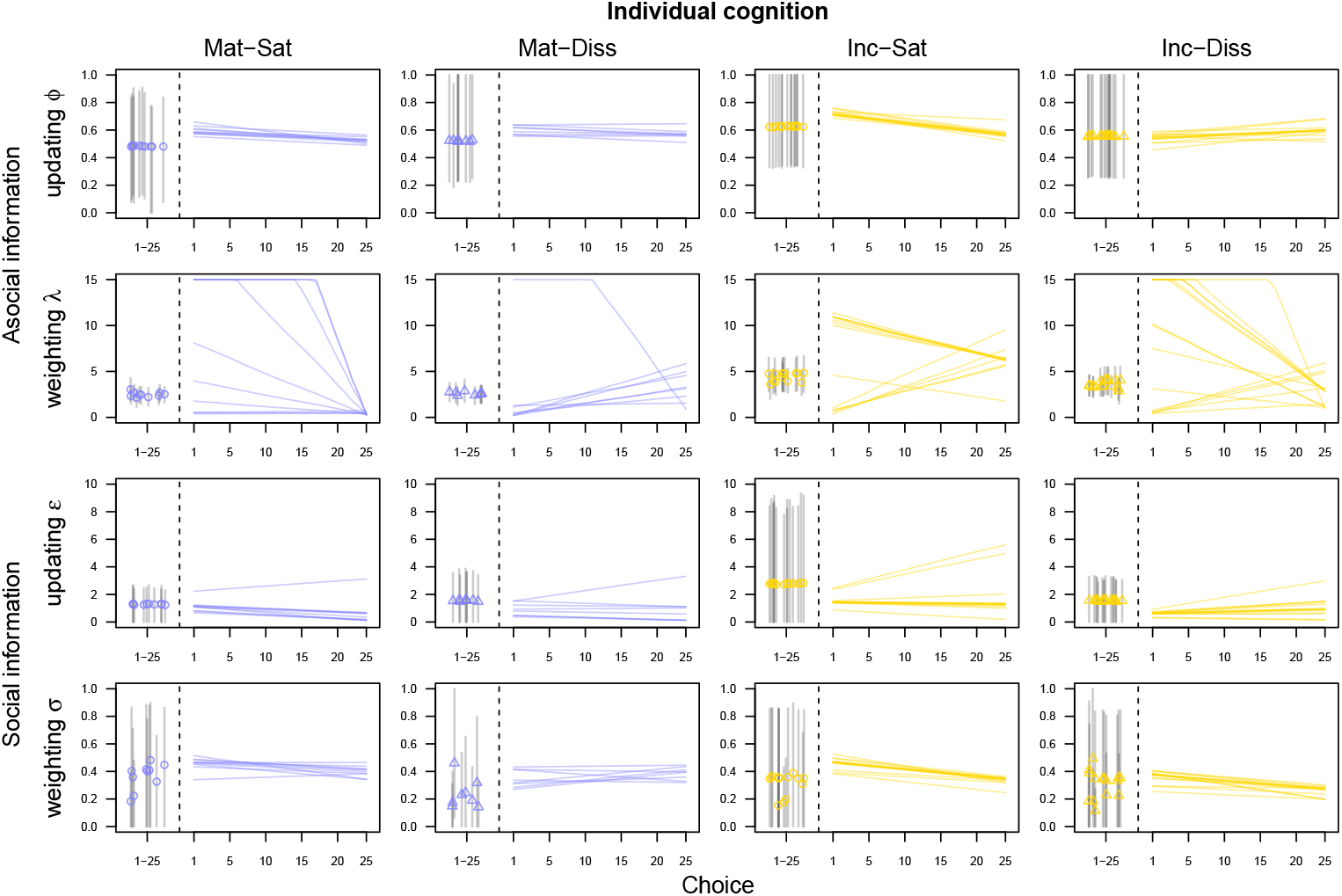
Individual-level cognition results. Latent individual-level (a)social learning parameters *ϕ, λ, ϵ*, and *σ* by male, treatment, and EWA model. Left panels display EWA baseline model estimates: open circles or triangles represent posterior means for each male, with black lines indicating 89% highest posterior density intervals. Right panels display EWA monotonic model estimates: each line shows one male’s posterior mean trajectory across 25 choices.

**Figure S4.**
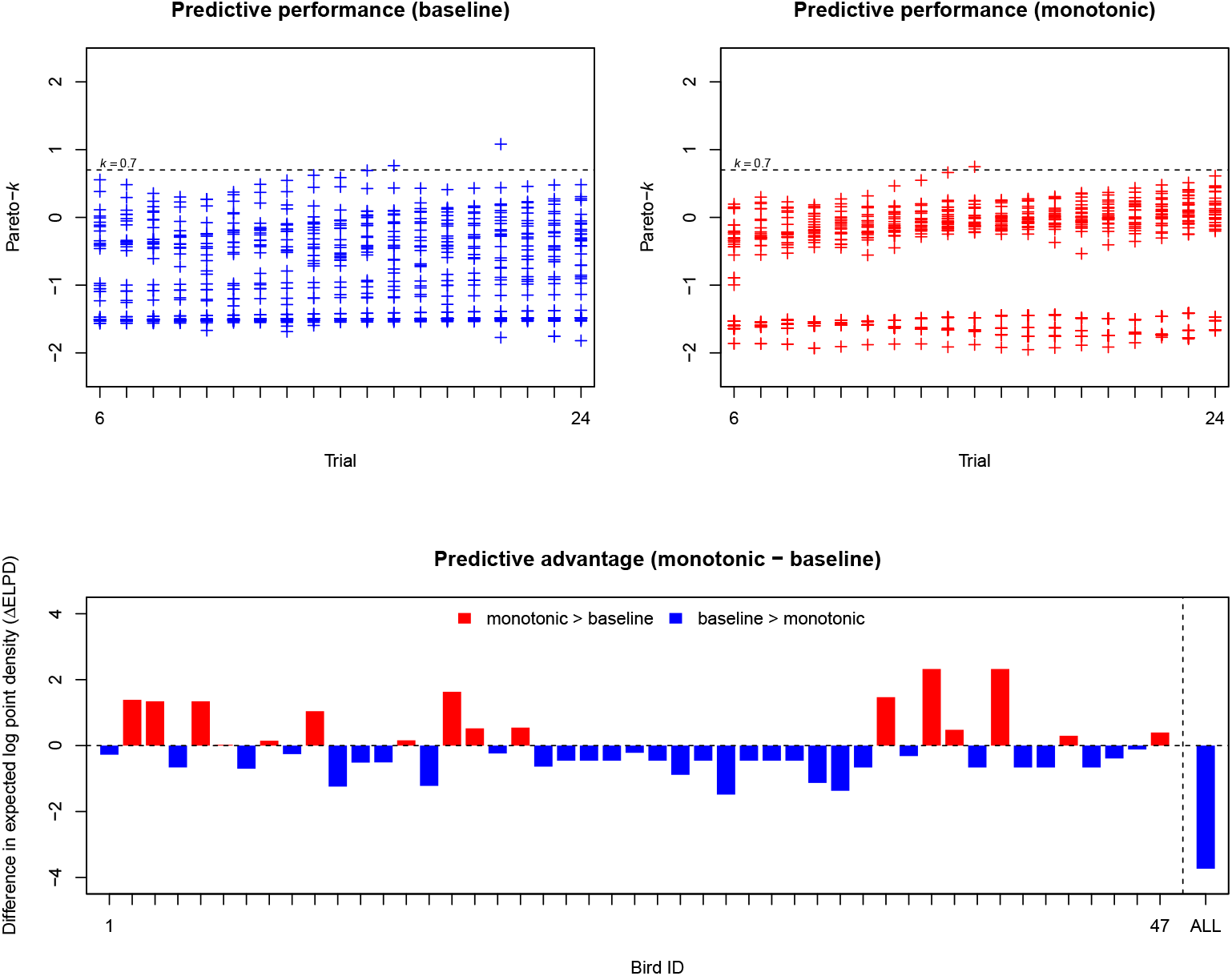
Model predictive performance. Top row: shows that each model’s Pareto-*k* values—predictive certainty indicators—were almost all below the *k* = 0.7 threshold (three exceptions), above which estimates became unstable and required refitting (twice for baseline, once for monotonic). These few refits indicate most trial-wise predictions were stable and the approximation held across individuals. Bottom row: comparison of ELPD values—predictive advantage measures—per individual and over “ALL” individuals (red = monotonic advantaged, blue = baseline advantaged). Overall, both models show similar performance (ELPD: baseline =− 266.79, monotonic =− 270.53), with a small baseline advantage (ΔELPD = − 3.73). Thus, both models made reliable predictions, supporting confidence in their outputs, with a modest baseline advantage. For an accessible overview of this forecasting method, see: https://mc-stan.org/loo/articles/loo2-lfo.html.

**Figure S5.**
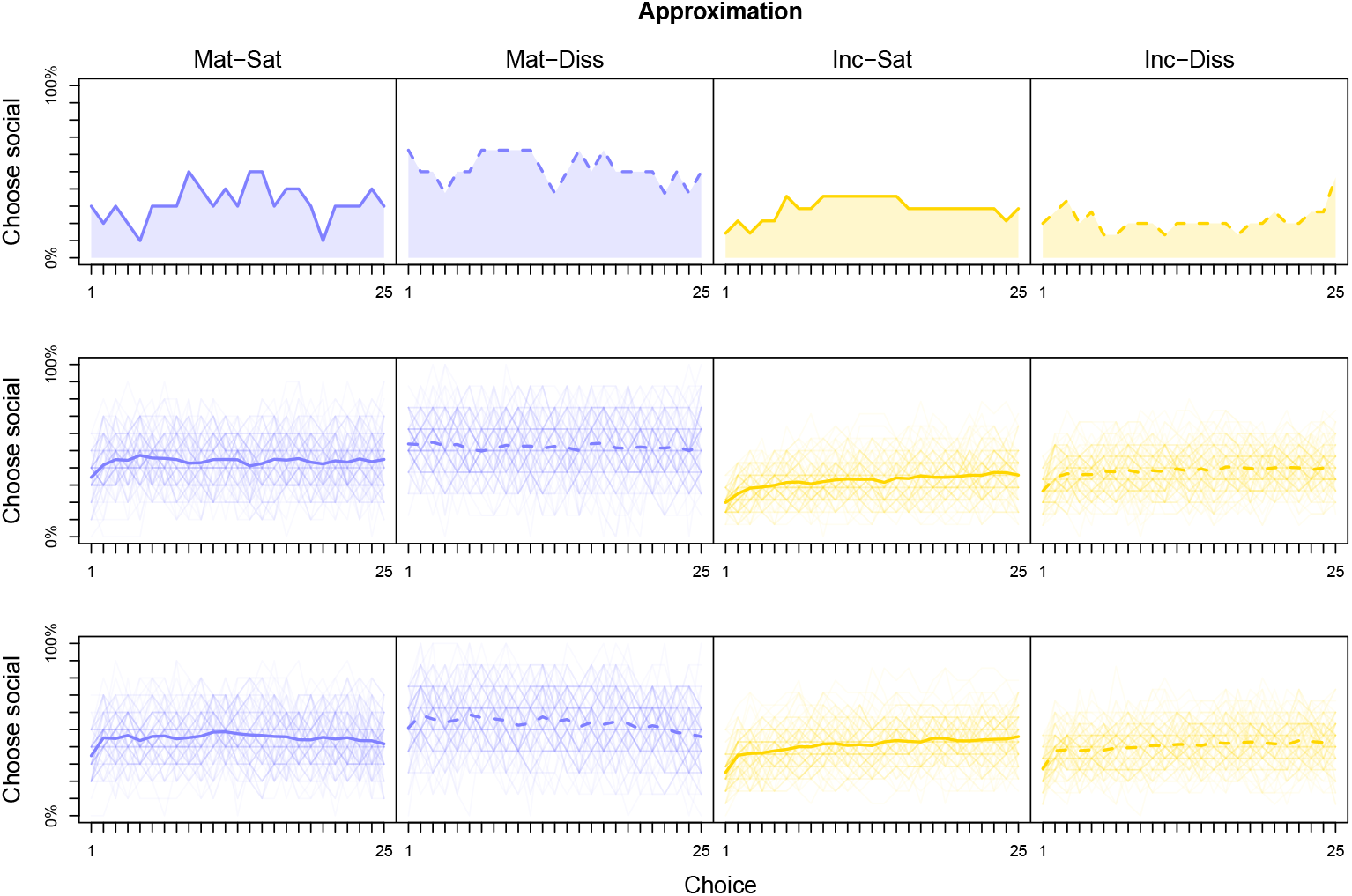
Forward simulation approximation results. Mean percentage social-material choice across 25 choices by treatment for: (top row) real birds; (middle row) ‘choosers’ generated from the posterior of the EWA baseline model; and (bottom row) ‘choosers’ generated from the posterior of the EWA monotonic model. Thick solid and dashed lines indicate the overall mean percentage of social material use across choices. Thin lines in the middle and bottom rows show the mean percentage social material use across choices for each simulation (*n* = 100). Simulations were matched to the sample size of each treatment. Both models tend to overestimate social material choice compared to the behaviour of real birds, except in the Mat-Diss treatment. However, ‘choosers’ generated from the EWA baseline model posterior slightly overshoot social material use to a lesser degree in the incubation treatments. See the main text for further discussion.

